# The maternal hormone in the male brain: sexually dimorphic distribution of prolactin signalling in the mouse brain

**DOI:** 10.1101/333161

**Authors:** Hugo Salais-López, Carmen Agustín-Pavón, Enrique Lanuza, Fernando Martínez-García

**Affiliations:** Unitat Predepartamental de Medicina, Facultat de Ciències de la Salut, Universitat Jaume I, Castelló de la Plana, Spain; Departament de Biologia Cel·lular i de Biologia Funcional, Facultat de Ciències Biològiques, Universitat de València, València, Spain

## Abstract

Research of the central actions of prolactin is virtually restricted to females, but this hormone has also documented roles in male physiology and behaviour. Here, we provide the first description of the pattern of prolactin-derived signalling in the male mouse brain, employing the immunostaining of phosphorylated signal transducer and activator of transcription 5 (pSTAT5) after exogenous prolactin administration. Next, we explore possible sexually dimorphic differences by comparing pSTAT5 immunoreactivity in prolactin-supplemented males and females. We also assess the role of testosterone in the regulation of central prolactin signalling in males by comparing intact with castrated prolactin-supplemented males.

Prolactin-supplemented males displayed a widespread pattern of pSTAT5 immunoreactivity, restricted to brain centres showing expression of the prolactin receptor. Immunoreactivity for pSTAT5 was present in several nuclei of the preoptic, anterior and tuberal hypothalamus, as well as in the septofimbrial nucleus or posterodorsal medial amygdala of the telencephalon. Conversely, non-supplemented control males were virtually devoid of pSTAT5-immunoreactivity, suggesting that central prolactin actions in males are limited to situations concurrent with substantial hypophyseal prolactin release (e.g. stress or mating). Furthermore, comparison of prolactin-supplemented males and females revealed a significant, female-biased sexual dimorphism, supporting the view that prolactin has a preeminent role in female physiology and behaviour. Finally, in males, castration significantly reduced pSTAT5 immunoreactivity in some structures, including the paraventricular and ventromedial hypothalamic nuclei and the septofimbrial region, thus indicating a region-specific regulatory role of testosterone over central prolactin signalling.

## INTRODUCTION

Prolactin (PRL) is a polypeptide hormone produced at the adenohypophysis, best-known for its role in the development of the mammary gland and milk production (1). To date, PRL has been attributed more than 300 different biological actions (2) in reproduction, homeostasis, angiogenesis or immunity, among others (3). To fulfil this array of functions, PRL has different target tissues, including the mammary gland, the uterus and other peripheral organs (3) and, importantly, the brain (4,5). In the brain, PRL is mainly involved in the regulation of a number of behaviours and physiological processes related to female reproduction and lactation (6). Indeed, PRL has been described as the “maternal hormone”, a pleiotropic hormone responsible for adapting every important aspect of female physiology and behaviour to the demands of motherhood (7).

The key role of prolactin in maternal physiology has traditionally biased neuroendocrinological studies towards its functions in the brain of females, whereas the actions of this hormone in the male brain have drawn less attention. For instance, the distribution of the PRL receptor (PRLR) has been reported for the female (8,9) but not the male rodent brain. One of the main reasons for this bias relates to the fact that male rodent models display low levels of circulating PRL under standard conditions (10), reflecting a limited access and functionality of PRL in the brain. Still, several studies do report acute rises in systemic PRL secretion in males associated to certain physiological conditions, such as the stress response (11) or sexual behaviour (12).

Regarding the latter, PRL is known to be involved in the control of male copulatory behaviour in rats (13), mice (14), other rodent models (15) and humans (16). Thus, PRL is released acutely with ejaculation (12) and it is proposed to intervene in the satiation following copulation(17) leading to the refractory period. In fact, chronic hyperprolactinaemia suppresses copulatory behaviour in animal models(18). In men, hyperprolactinaemia associated to several clinical conditions (19) (including antipsychotic treatments (20)), yields sexual dysfunction and other side-effects (21). The inhibitory role of PRL on sexual function is proposed to have an important central component (22,23), but the exact regions and mechanisms through which PRL operates in the male brain are still poorly understood.

In this light, the main aim of this work is to provide a comprehensive description of the distribution of cells sensitive to PRL in the brain of males. To do so, we have supplemented males with a standard dose of exogenous PRL and revealed PRL-responsive cells in the brain by means of the immunohistochemical detection of the phosphorylated form of signal transducer and activator of transcription 5 (pSTAT5). Phosphorylation of STAT5 is a key event in the Jak/STAT pathway, the major signalling cascade of the PRL receptor (3). In contrast to other methods, e.g. in situ hybridization of the PRL receptor (PRLR) mRNA (or its immunodetection), pSTAT5 immunohistochemistry in PRL-supplemented animals reveals those cells that are not just expressing PRL receptors or specific elements of their signalling cascade, but those that are fully responsive to PRL. Thus, pSTAT5 immunodetection can be considered a reliable method to assess central PRL responsiveness (4,5)

The second goal of this work is to compare the pattern of central PRL-derived signalling between male and female mice, in search for intersexual differences. For this purpose, we compare qualitatively and quantitatively the patterns of our PRL-supplemented males with ovariectomised, steroid-primed females treated with an equivalent dose of PRL. In addition, a direct comparison of the distribution of pSTAT5 in the brain of males and females (where most functional studies on the matter have been conducted) might also be useful for the functional contextualization of PRL input to the male brain.

In accordance with the involvement of PRL in reproductive physiology and behaviour, gonadal steroids are important regulators of PRL function in the brain. In female rodents, estradiol and progesterone have been shown to target most PRL-sensitive neurons in the brain (24) and to regulate PRL signalling at different levels (25–27). On the other hand, testosterone has been documented to inhibit PRL receptor expression in the pituitary gland of male rats and pituitary PRL release in male rats and mice (28–30). Beyond these findings, to our knowledge there is currently no available evidence supporting a regulatory role of testosterone in brain PRL signalling. Hence, in this work we also explore the putative contribution of testosterone in shaping the patterns of PRL responsiveness in the male brain. To this end, we analyse the levels of pSTAT5-ir in the main PRL-responsive regions of the male mouse brain after permanent testosterone withdrawal through castration, as compared to intact male mice, both supplemented with PRL.

By characterizing the patterns of PRL signalling in the male brain, assessing its sexual dimorphism and clarifying their dependence on testosterone, we intend to shed light on the roles of PRL in male behaviour and physiology, as well as on the neural substrates for PRL-dependent, male-specific behaviours.

## MATERIAL AND METHODS

### Animals

For the present study, we used 15 male and 6 female mice of the CD1 strain (Charles River Laboratories, France), aging between 8 and 24 weeks. These animals were housed in polypropylene plastic cages under controlled temperature (24 ± 2 <u>°</u>C) and lighting conditions (12h:12h; lights ON at 8 am), with *ad libitum* access to food and water. Intact males were housed individually (to avoid inter-male aggression) whereas castrated males and females were group-housed (4 to 6 animals per group, to avoid isolation-derived stress). Animals were treated throughout according to the European Union Council Directive 2010/63/EU (6106/1/10 REV1) and procedures were approved by the Committee of Ethics on Animal Experimentation of the University of Valencia (protocol number: 2015/VSC/PEA/00055), where the experiments were performed.

### Experimental Design

Male mice were randomly assigned to three experimental groups: (1) Male Control (n=3); (2) Male+PRL (n=6) and (3) Castrated+PRL (n=6). Animals of Male Control and Male+PRL groups were left gonadally intact, whereas males from Castrated+PRL group underwent orchidectomy. Following the procedure by Brown and collaborators (5,31), Male+PRL and Castrated+PRL groups received an exogenous PRL supplementation 45 minutes prior to perfusion.

In order to compare central PRL sensitivity between males and females, we also included in our study a group of ovariectomised, steroid-primed and PRL-supplemented females, which belonged to a sample of a previously published set of experiments (4). The steroid-replacement schedule conducted on these females intended to emulate the proestrus-estrus phase of the estrous cycle (32). Given the considerable variability of pSTAT5-ir in the brain of freely-cycling females, likely due to gonadal steroid influence on PRL signalling (4), this group provided a more stable comparison than freely-cycling females and a more accurate comparison than estrous cycle screening.

Data from these four groups of animals (Male Control; Male+PRL; Castrated+PRL and Female+PRL) allowed us to: a) characterize the pattern of PRL-derived signalling in the male brain by analysing the distribution of pSTAT5-ir in group Male+PRL; b) analyse the effect of PRL supplementation on pSTAT5-ir distribution by comparing the Male Control with Male+PRL groups; c) explore sexual dimorphism in the central sensitivity for PRL by comparing Male+PRL with Female+PRL groups; and finally, d) assess the modulatory role of testosterone on PRL signalling in the male brain through the comparison of the Male+PRL and the Castrated+PRL groups.

### Ovariectomy and Orchidectomy

Males of Castrated+PRL group were orchidectomised at approximately 12 weeks of age. Briefly, animals were deeply anaesthesised with i.p. ketamine (Imalgene 500, Merial, Toulouse, France, 75mg/kg) and medetomidine (Domtor 1mg/ml, Esteve, Barcelona, Spain, 1 mg/kg) and surgery was performed via a single midline incision on the scrotal sac. After surgery, i.p. atipamezol hydrochloride (Antisedan, Pfizer, New York, USA, 1 mg/kg) was administered to reverse anaesthesia and s.c. butorphanol tartrate 1% (Torbugesic, Pfizer, New York, USA, 20 µl) was regularly administered through the postsurgical period for pain control. Castrated males had 2-3 weeks of recovery under undisturbed housing conditions before the beginning of the experiments.

Experimental females underwent ovariectomy 9-10 weeks after birth under identical pharmacological conditions as explained for males (4). Surgery was conducted through two incisions on both sides of the back of the animal. Ovariectomized females had at least seven days of recovery after surgery.

### Hormone Treatments

Experimental females underwent a steroid replacement schedule (4) consisting of a sustained treatment with 17-β estradiol and an acute treatment with progesterone. In accordance with the experimental induction of estrus (32), estradiol was administered on a slow-release profile during 7 days, by means of the subcutaneous placement of silastic tubing implants (Dow Corning Corporation, Midland, MI, USA) filled with 20 µg/ml β-estradiol (Sigma, St Louis, MO, USA) diluted in sunflower oil. Silastic tubing had an inner diameter of 1.67 mm and an outer diameter of 2.41 mm, and implants were cut to a length of 20 mm. Implants were inserted subcutaneously on the lumbar region under isoflurane anaesthesia (Isoflo, Esteve Veterinaria, Barcelona, Spain). Animals were also administered a subcutaneous dose of 20 µl butorphanol tartrate 1% (Torbugesic, Pfizer, New York) during implant placement surgery, for pain control. By contrast, progesterone (Sigma, St Louis, MO, USA) was administered to females acutely in a 500 µg subcutaneous injection, diluted in sunflower oil, in the morning of the seventh day of estradiol treatment, specifically 3 hours and 45 minutes prior to perfusion. In order to make males and females directly comparable for the analysis of sexual dimorphism, males of the Male+PRL group received vehicle treatment parallel to that of females (implants with sunflower oil, plus s.c. oil injection 3h 45 minutes prior to perfusion).

Regarding PRL supplementation, except for the Male Control group (which received vehicle injection), all experimental groups were administered an acute 5 mg/kg i.p. dose of ovine PRL (Sigma, St Louis, MO, USA) 45 minutes before perfusion. The dose and timing of PRL administration rendered homogenous, supraphysiological circulating levels of PRL and ensured that the peak of STAT5 phosphorylation in response to a PRL challenge coincided with perfusion (5). This is intended to reveal the maximal potential response patterns among experimental animals and also to allow direct comparison between experimental groups, regardless of sex (male vs female) or physiological condition (intact vs castrated males).

### Tissue Collection and Histological Processing

Animals received an overdose of sodium pentobarbital (Vetoquinol, Madrid, Spain) and were transcardially perfused with 4% paraformaldehyde in 0.1M phosphate buffer (PB), pH 7.4. Brains were carefully extracted and post-fixed overnight through immersion in the same fixative, then cryoprotected by immersion in 30% sucrose in 0.01 M PB until they sank (2-3 days). Using a freezing microtome (Microm HM-450, Walldorf, Germany) brains were sectioned in five parallel series of 40µm thick coronal sections. Series were stored in PB-30% sucrose at -20<u>°</u>C until their use.

### Immunohistochemistry for pSTAT5

Immunohistochemistry was conducted in free-floating sections under light shaking at room temperature (25<u>°</u>C) unless otherwise stated. Immunohistochemistry protocol was adapted from Brown et al.(5,31). Tissue sections were thoroughly rinsed between stages for at least three 10-min washes in TRIS-buffered saline, 0.05M, pH 7.6 (TBS). After thawing, sections underwent an initial antigen retrieval step, consisting in two sequential 6-minute incubations in 0.01 M TRIS buffer (TB), pH 10 at 85<u>°</u>C, and brought quickly to room temperature in between. Tissue was then incubated in: a) 1% hydrogen peroxide (H_2_O_2_) for 30 minutes, for endogenous peroxidase inhibition; b) 2% BSA, 2% goat serum and 0.3% Triton X-100 in TBS for 1h, in order to block unspecific labelling; c) rabbit monoclonal anti-pSTAT5 primary antibody (Phospho-Stat5 Tyr694 (D47E7), Catalog number #4322, Cell Signaling Technology, Danvers, MA) diluted 1:500 in TBS plus Triton X-100 0.1% for 72 h at 4<u>°</u>C; d) biotinylated goat anti-rabbit IgG (Vector Laboratories, Peterborough, UK; RRID AB_2313606) 1:200 in TBS for 90 minutes; and e) avidin-biotin-peroxidase complex (ABC Elite kit; Vector Laboratories; RRID AB_2336819) in TBS for 90 minutes. Peroxidase label was developed using 0.005% 3-3’-diaminobenzidine (Sigma) and 0.01% H_2_O_2_ in TB pH 7.6 for about 15 minutes, obtaining thereby a brown nuclear staining. Sections were rinsed in TB and mounted onto gelatinized slides, dehydrated in graded ethanol, cleared with xylene and coverslipped with Entellan.

The specificity of the anti-pSTAT5 primary antibody was demonstrated by means of an antibody competition assay (Fig S1).

### Analysis of pSTAT5 Immunoreactivity

The patterns of pSTAT5-ir distribution were mapped for the brain of intact males (Male+PRL) and females (Female+PRL) and illustrated semiquantitatively in camera lucida drawings of selected brain sections (Fig 1). In addition, a quantitative assessment of the density of cells showing pSTAT5 immunoreactivity (pSTAT5-ir) in representative brain sites was performed for PRL-treated females and males, both intact and castrated. To do so, we first selected frames of the chosen nuclei using the stereotaxic atlas of Paxinos and Franklin (2004). Then, we obtained photomicrographs of these frames in both hemispheres using a digital camera (Leica DFC495) attached to a microscope Leitz DMRB (Leica AG, Germany). Thereafter, images were processed and analysed using ImageJ. Briefly, we subtracted background light and converted the RGB colour image to greyscale by selecting the green channel. Then, we binarised the greyscale image setting the 75% of the mode of the histogram as a threshold, thus including every pixel below this threshold as positively labelled. Using the ImageJ commands “fill holes”, “open” (3 iterations) and “watershed”, we filtered small particles (noise or fragmented nuclei) and separated fused objects. The resulting particles were additionally filtered by area (larger than 70 µm^2^, corresponding to an approximate diameter of 9.4 µm) and finally counted automatically. We calculated the mean (interhemispheric) density of pSTAT5-imunoreactive cell nuclei for each specimen by dividing the counts for both hemispheres by the area of both frames.

**Figure 1.**
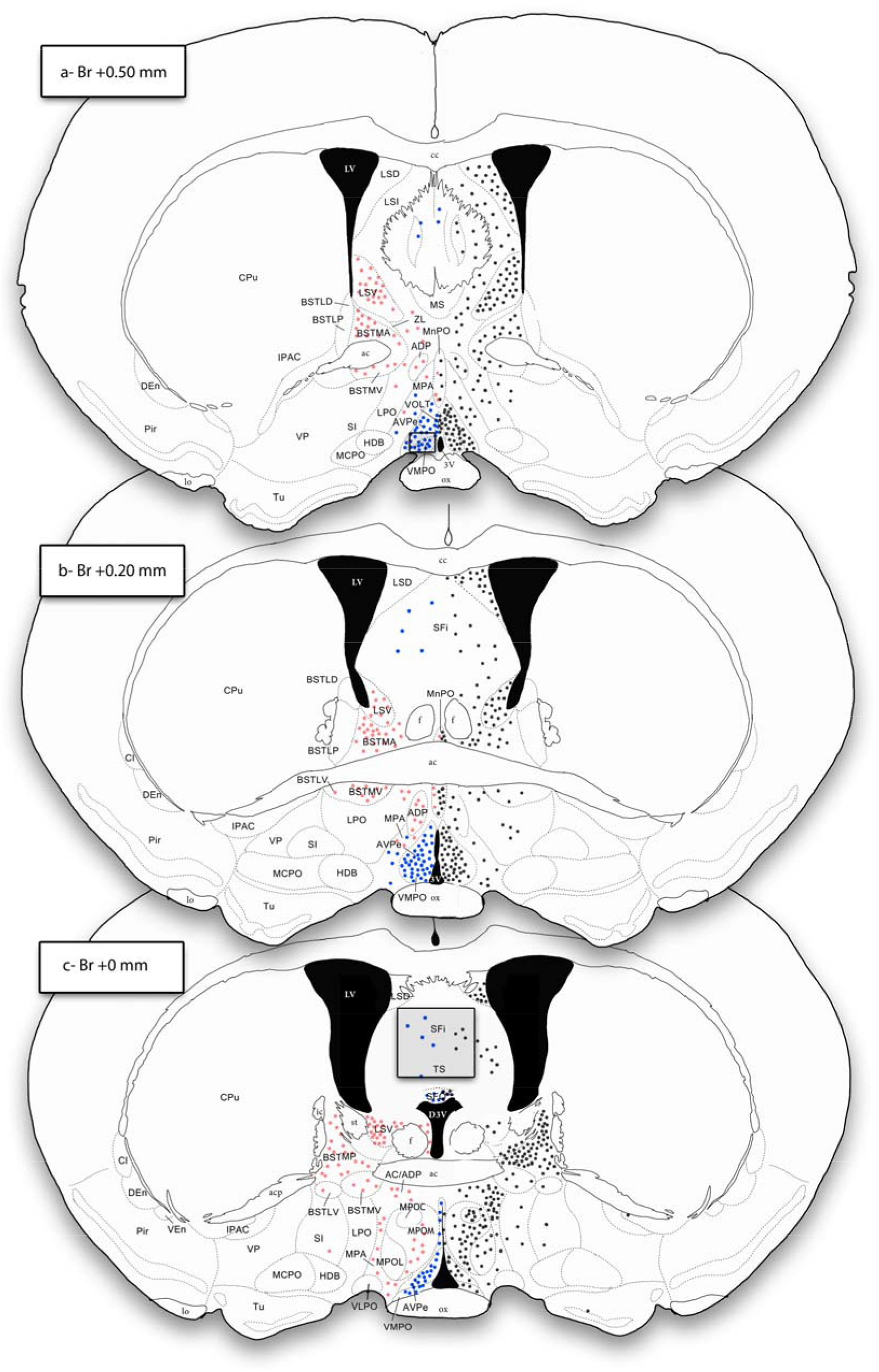
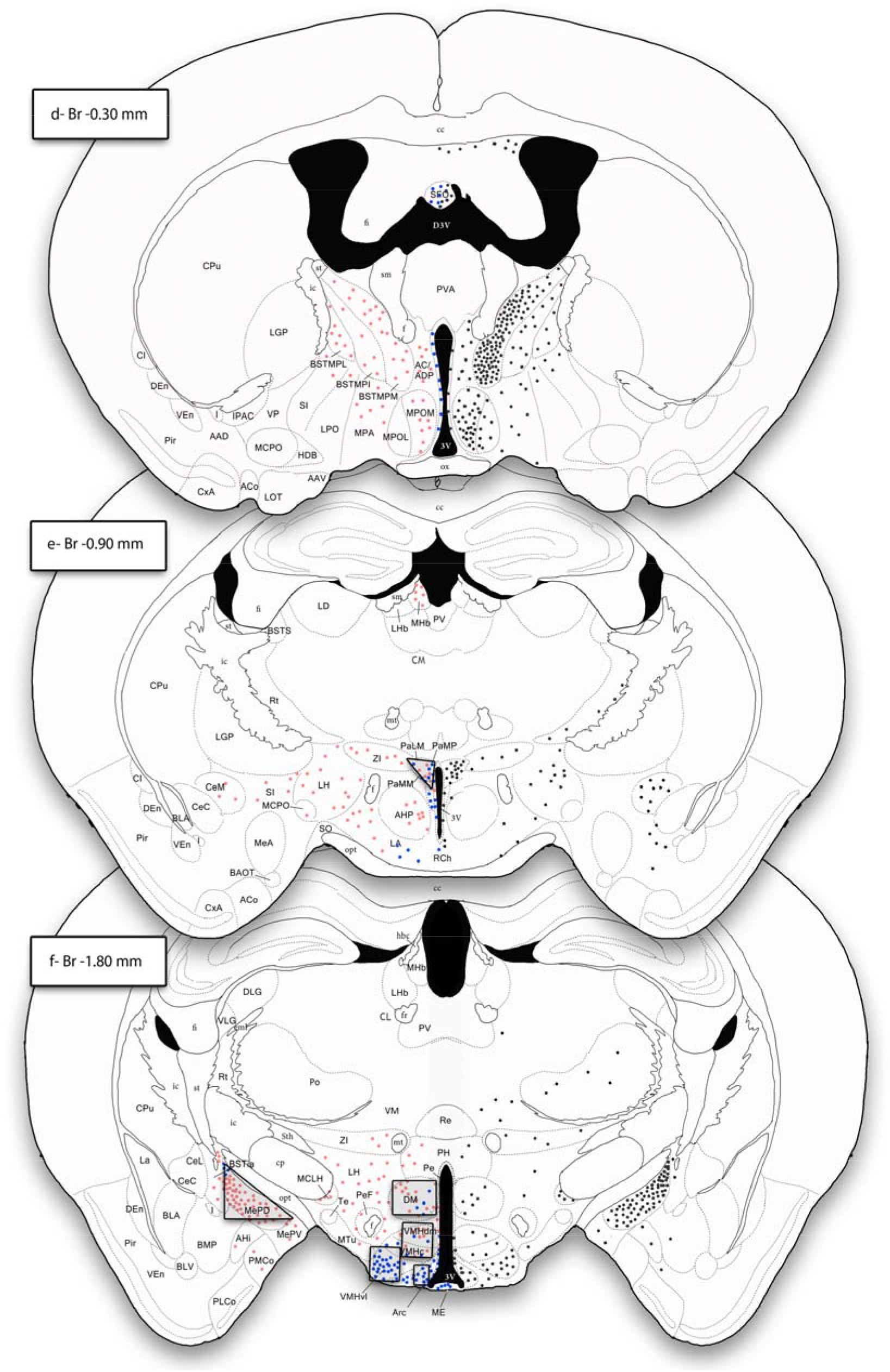
Mapping of pSTAT5 immunoreactivity and prolactin receptor expression in the brains of male and female mice. Semi-schematic camera lucida drawings of coronal sections of the mouse brain showing the distribution patterns of pSTAT5-ir in female and male mice (left side) and the pattern of PRLR expression in the brain of an adult male mouse specimen (right side), as determined by PRLR expression data obtained from the Allen Brain Institute (© 2004 Allen Institute for Brain Science. Allen Mouse Brain Atlas. Available from: mouse.brain-map.org experiment 72340223, http://mouse.brain-map.org/experiment/show/72340223). In the left side, pink dots represent pSTAT5 expression exclusive of ovariectomized, steroid-primed females, whereas dark blue dots encode overlapping expression of pSTAT5-ir in both the female and male specimens. Black dots in the right illustrate PRLR expression in the male mouse brain. Each dot represents approximately 4 cell nuclei labelled for pSTAT5 or equivalent number of cells expressing the PRLR. Shaded areas indicate the counting frames designed for the AVPe/VMPO region (Fig 1a), the Pa (Fig 1e), and the Arc, VMHvl, VMHc and VMHdm, DM and MePD (all frames in Fig 1f), as part of the quantitative analysis performed in this work (see below). Approximate distance to bregma is indicated for each section.

### Statistical Analysis

We performed statistical analysis of the resulting data on the SPSS software package. After checking for normality (Kolmogorov-Smirnov test with Lilliefors’ correction) and homogeneity of variances (Levene’s test), we conducted three different statistical comparisons. First, we compared levels of pSTAT5-ir density between intact males (Males + PRL) and ovariectomised, steroid-primed females (Female+PRL), in search for dimorphic differences. Then, we checked for the effect of orchidectomy on male pSTAT5-ir levels by comparing intact (Male+PRL) and castrated (Castrated+PRL) males. For both analyses, the samples fulfilling the criteria for a parametric analysis were subject to an independent t-test, whereas the samples of the remaining nuclei were subject to a non-parametric Mann-Whitney test. For each statistical test, we applied a significance level of 0.05.

## RESULTS

The main focus of this section is to characterise the central patterns of pSTAT5-ir found in male mice in the presence of high levels of circulating PRL (Male+PRL group), as an indicator of PRL-derived signalling occurring in the male brain. Likewise, we compare the distribution of pSTAT5-ir between PRL-supplemented males (Male+PRL group) and non-supplemented males (Male Control group). Then, we analyse intersexual differences in PRL-derived signalling in the mouse brain. For this purpose, we compare the patterns of pSTAT5-ir in the male (Male+PRL group) and female (Female+PRL group) brains and assess the density of pSTAT5-ir in a number of brain regions displaying PRL-derived signalling in both males and females. Finally, we explore the effect of testosterone withdrawal in the PRL-derived signalling of the male brain by comparing pSTAT5-ir levels between intact and castrated male mice. For the following anatomical descriptions, we adhere to the neuroanatomical terminology proposed by Paxinos and Franklin(33) (see also list of abbreviations above).

### A. Patterns of pSTAT5 immunoreactivity in the male mouse brain: effect of PRL supplementation

Immunohistochemistry for pSTAT5 produced a defined staining in the examined brain tissue, which was restricted to the cell nucleus (see Fig 2), as shown in our previous work (4). Labelling was suppressed by pre-incubation of the antibody with the phosphorylated peptide used for immunization, but not when the antibody was incubated with the equivalent, non-phosphorylated peptide (Fig S1). Therefore, we safely assume that, in our immunohistochemical procedure, the antibody is detecting specifically pSTAT5.

**Figure 2.**
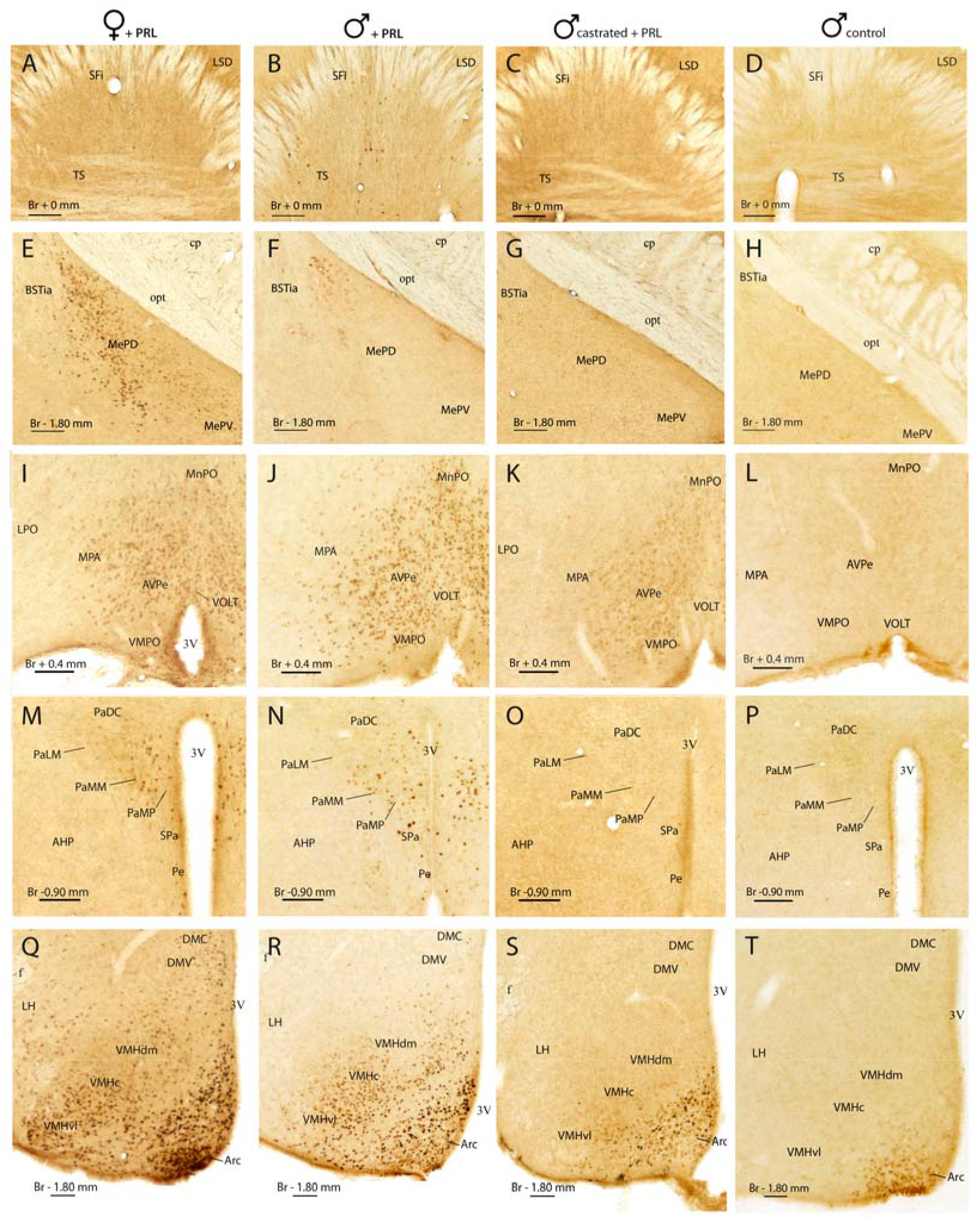
Representative examples of pSTAT5 immunoreactivity in the brain of females, intact males and castrated male mice. Photomicrographs illustrating pSTAT5 labelling in representative brain sections in an ovariectomized, steroid-primed female supplemented with PRL (I), an intact, PRL-supplemented male (II), a castrated, PRL supplemented male (III) and a intact control male lacking PRL supplementation (IV). Sections correspond to the preoptic hypothalamus (A), the paraventricular region in the anterior hypothalamus (B), the tuberal hypothalamus (C), the caudal septum (D) and the medial posterior amygdala (E). The approximate distance to bregma is indicated in each section. Scale bars correspond to 100 µm.

Figure 1 shows the distribution of pSTAT5-ir observed in PRL-supplemented males (blue dots on the left side of the brain), as well as the the distribution of cells expressing PRL receptor (PRLR) in the male mouse brain (Fig 1, black dots on the right side of the brain), as reported by the Allen Brain Institute (Allen Mouse Brain Atlas, experiments 1268 and 72340223, Fig S2). Figure 2 shows photomicrographs illustrating pSTAT5-ir in selected brain nuclei for all of our experimental groups.

Males supplemented with PRL (Male+PRL) showed moderate-to-dense pSTAT5-ir in several nuclei of the hypothalamus and cerebral hemispheres (Fig 2, left column), as well as in the choroid plexus (not shown). In the telencephalon, pSTAT5-ir was mainly observed in the septum, specifically in the area comprising the septofimbrial nucleus (SFi) and the triangular septal nucleus (TS) (Figs 1B and 1C). In this region, the subfornical organ (SFO) also showed some immunoreactive cells. In addition, the posterodorsal nucleus of the medial amygdala (MePD) displayed scarce pSTAT5-ir, with few immunolabelled cells restricted to the upper corner of the nucleus, in close contact to the intramygdaloid BST (BSTia, see Figs 1F and 2F). Importantly, all these brain nuclei display PRLR mRNA expression in the male mouse (Fig 2 right side).

In the hypothalamus, pSTAT5-ir was present in the preoptic, anterior and tuberal regions. In the preoptic hypothalamus (Figs 1A-D and 2J), pSTAT5-ir cells were concentrated in the juxtaventricular nuclei, namely the vascular organ of the *lamina terminalis* (VOLT) and the anteroventral periventricular (AVPe), the ventromedial preoptic (VMPO) and the Pe nuclei. In addition, the adjoining MPA displayed a few immunolabelled cell nuclei.

In the anterior hypothalamus, pSTAT5-ir was once again restricted to juxtaventricular structures, the paraventricular nucleus (Pa) and the Pe (Figs 1E and 2N), as well as the retrochiasmatic area (RCh, Fig 1E). In the Pa, labelling was restricted to the medial aspect of the nucleus, mainly to the ventral (PaV), medial magnocellular (PaMM) and medial parvocellular (PaMP) subdivisions, with very few labelling observed in the lateral aspect of the nucleus (PaLM) or the subparaventricular nucleus (SPa). In the tuberal hypothalamus (Figs 1F and 2R), the arcuate nucleus (Arc) displayed abundant pSTAT5-ir, with some immunolabelled cells displaced into the median eminence (ME). In some males, the dorsomedial nucleus (DM) displayed few scattered immunolabelled cells. Finally, pSTAT5-ir appeared in the ventromedial hypothalamic nucleus (VMH), with higher levels in the ventrolateral subdivision (VMHvl) and sparse labelling in the rest (VMHc and VMHdm).

In contrast to the Male+PRL group, non-supplemented males (Male Control, Fig 2, rightmost column), were virtually devoid of pSTAT5-ir, except for some labelled cells in the ventral aspect of the arcuate nucleus (Arc, Fig 2T) and the anterior aspect of the periventricular nucleus (Pe, not shown).

### B. Comparative analysis of pSTAT5-ir the male and female mouse brain

In comparison to males, ovariectomised, steroid-primed females supplemented with PRL (Female+PRL) showed a more extensive distribution of pSTAT5-ir (Fig 1, Fig 2 middle-right column), which matches previous descriptions conducted on freely-cycling female mice (4). To summarize, labelling in our experimental females was widespread in the basal telencephalon and hypothalamus, but was also found within thalamic, midbrain and brainstem structures. To allow a direct comparison between sexes, Figure 2 shows the distribution of pSTAT5-ir in equivalent brain sections of male and female mice. In addition, in the semischematic drawings of brain sections depicted in Fig 1, pSTAT5-ir labelling that was common to males and females is shown as blue spots, whereas labelling exclusively present in females is illustrated as red spots. Importantly, there was not a single brain site where labelling was exclusively found in males. In other words, after administration of equivalent doses of exogenous PRL, males show pSTAT5-ir in a reduced number of brain centres, whereas females show immunoreactive cells in the same nuclei, plus a set of additional brain centres. This represents a clear case of sexual dimorphism in favour of females.

We also checked for quantitative sexual dimorphism in PRL-derived signalling, by analysing the density of pSTAT5-ir in the main nuclei showing pSTAT5-ir in PRL-supplemented intact males and steroid-primed ovariectomised females. We designed counting frames for the following brain nuclei: the AVPe/VMPO region, the Pa, the Arc, the DM, the VMHvl, the area comprising the central and dorsomedial VMH (VMHc and VMHdm, respectively), the MePD and the SFi (Fig 1). These results are summarized in Figure 3. Females showed a general trend towards higher levels of pSTAT5-ir than males. This effect reached statistical significance in the Arc (t(10)=3.043, p=0.012), the DM (U=32.0; p=0.026) and the MePD (U=30.0; p=0.004), although the AVPe/VMPO also showed an almost significant trend towards higher pSTAT5-ir density in females (t(9)=2.183; p=0.057).

**Figure 3.**
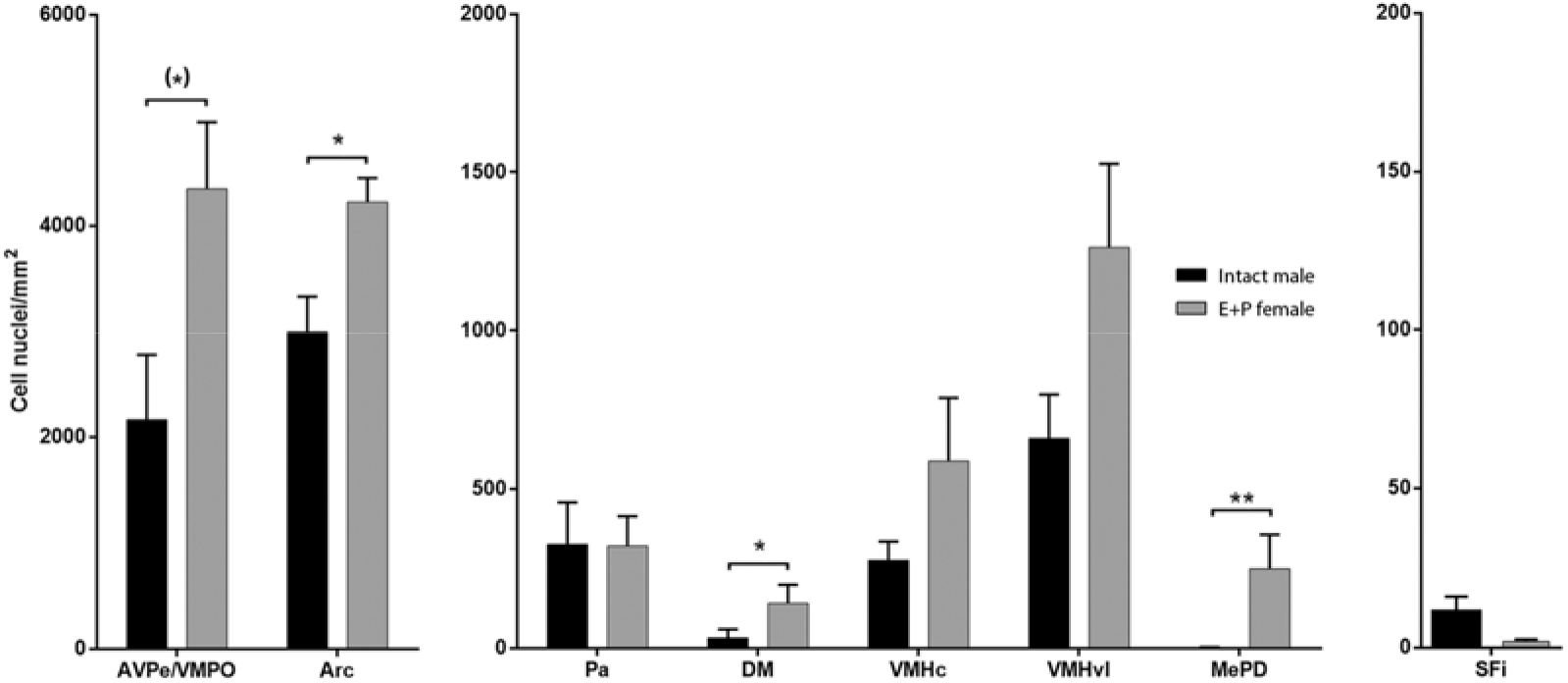
Quantitative analysis of pSTAT5 immunoreactivity in selected brain regions of female and male mice. Assessment of pSTAT5-ir density (pSTAT5-positive cell nuclei/mm^2^) in the major brain regions with expression of pSTAT5-ir in both male and female mice. Bar histograms show mean interhemispheric pSTAT5-ir density ± SEM in gonadally-intact, vehicle-treated males (Male+PRL; n=6; black) and ovariectomized females treated with estradiol and progesterone (Female+PRL; n=6; grey). Counting frames for each of the analyzed nuclei are included in Fig1. Statistical analysis was applied independently to each brain region (independent t-test for parametric data or Mann-Whitney test for non-parametric data, see Results). *P ≤ 0.05; **P ≤ 0.01; (*) P ≤ 0.06.

### C. Effect of testosterone withdrawal in PRL-derived signalling in the male mouse brain

Finally, we explored the putative role of testosterone in the regulation of PRL-derived signalling in the male brain. To do so, we compared pSTAT5-ir patterns between intact, PRL-supplemented males (Male+PRL; Fig 2, second column from left) and a sample of castrated, PRL-supplemented males (Castrated+PRL; Fig 2, third column). In qualitative terms, castrated and intact male mice displayed similar patterns of pSTAT5-ir, but castrated males showed apparently lower levels of pSTAT5-ir. Quantitative analysis of the labelling in both groups (using the same methodology and exploring the same subset of nuclei analysed in the previous section) allowed comparing pSTAT5-ir between gonadally intact and castrated males by means of independent t-tests for the Arc and AVPe and non-parametric Mann-Whitney tests for the rest of the analysed regions. The results of this statistical comparison reveal a significant effect of testosterone withdrawal in diminishing the levels of pSTAT5-ir in the Pa (U= 33.00; p=0.004), in both the ventrolateral and central/dorsomedial VMH (U= 30.00, p=0.015 for both frames) and in the SFi (U= 26.00, p=0. 05). In the remaining analysed nuclei, the density of pSTAT5-ir cells was similar in Male+PRL and Castrated+PRL animals, as no statistical differences were found (p>0.1) (Fig 4).

**Figure 4.**
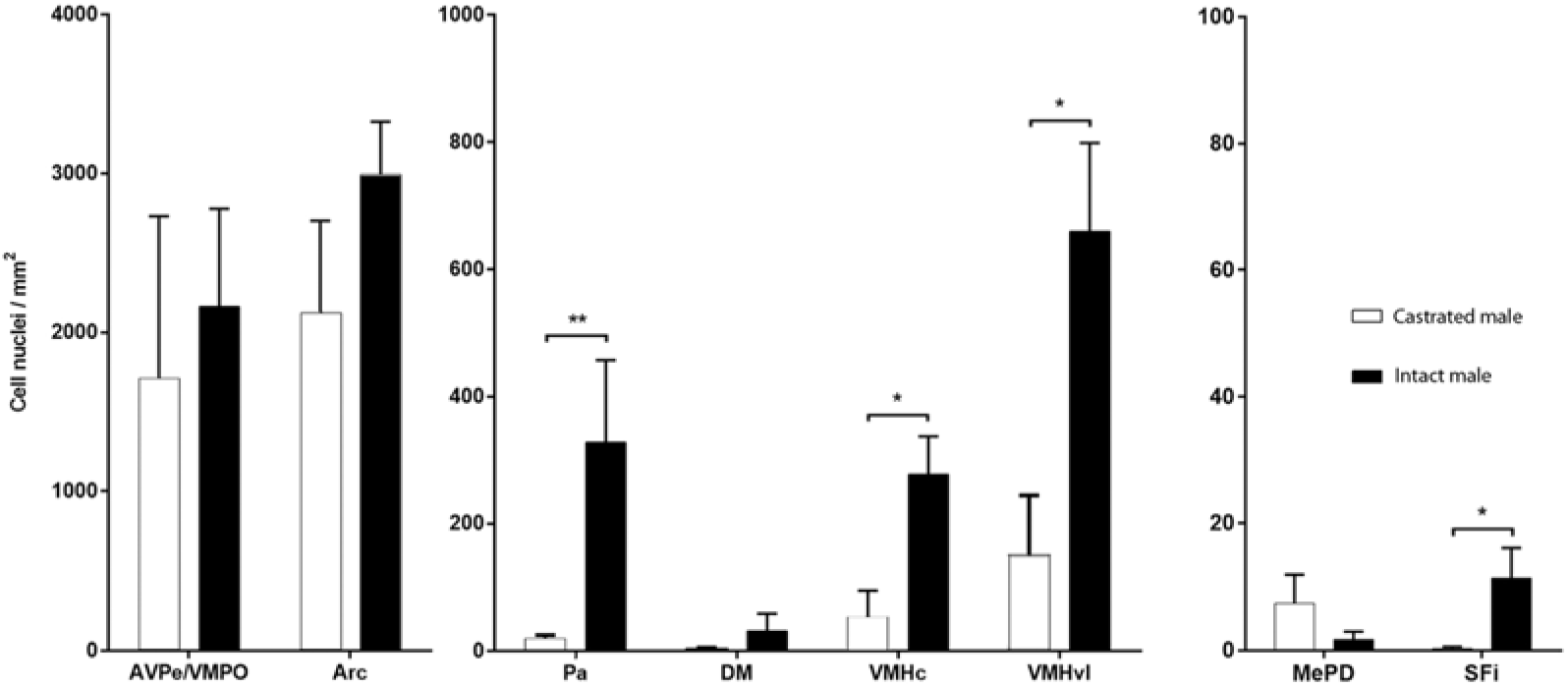
Effect of testosterone withdrawal in pSTAT5 immunoreactivity of the male mouse brain. Assessment of pSTAT5-ir density (pSTAT5-positive cell nuclei/mm^2^) in gonadally-intact male mice and castrated male mice within the major brain regions showing pSTAT5 expression in the male mouse brain. Bar histograms show mean interhemispheric pSTAT5-ir density ± SEM in intact, PRL-supplemented males (Males+PRL; n=6; black)and castrated, PRL-supplemented males (Castrated+PRL; n=6; white). Counting frames for each of the analyzed nuclei are enclosed in Fig 2. Statistical analysis was applied independently to each brain region (independent t-test for parametric data or Mann-Whitney test for non-parametric data). *P ≤ 0.05; **P ≤ 0.01.

## DISCUSSION

The present study examined, for the first time, the responsiveness of the brain of males to PRL. Specifically, we analysed the distribution and steroid regulation of PRL-derived signalling in the male mouse brain, by means of the immunohistochemical detection of pSTAT5. We also compared the pattern of pSTAT5 immunoreactivity in the brain of males and females, aiming at assessing its possible sexual dimorphism. In this section, we will first analyse the distribution of pSTAT5 in the brain of male mice, with or without exogenous PRL administration, and compare it to the pattern of expression of the PRLR, as shown in in situ hybridisation material provided by the Allen Brain Institute (see Fig 1 and Fig S2). Then, we will comment on the observed sexually dimorphic differences in central sensitivity to PRL. Next, we will discuss the role of testosterone as a putative regulator of central PRL signalling and on the mutual regulatory relationship of PRL and testosterone. We will conclude by reviewing how our findings contribute to elucidate the neuroanatomical substrate of some putative actions of PRL in the brain of males.

### Prolactin signalling in the male mouse brain, PRL supplementation and PRL receptor expression

Although the actions of PRL in the female mouse brain have received considerable attention due to the role of PRL in female reproduction and motherhood(34), this work is, to our knowledge, the first descriptive analysis of the distribution of PRL-derived signalling in the male mouse brain. In PRL-supplemented male mice (Male+PRL group), brain regions positively labelled for pSTAT5 were located mainly in the hypothalamus (the AVPe and VMPO, MPA, Pe, Pa, Arc, DM and VMH) and, to a lesser extent, in the telencephalon (SFi in the septum and MePD in the amygdala), with no additional pSTAT5-ir detected in any of the remaining brain divisions (Fig 1).

Importantly, this pattern of pSTAT5-ir was highly dependent on systemic supplementation with a high dose of exogenous PRL, whereas non-supplemented controls (Male Control group) were virtually devoid of pSTAT5-ir, except for some faintly labelled cells in the ventral Arc (Fig 2T) and in the anterior portion of the periventricular hypothalamus (not shown). These results suggest, in the first place, that the patterns of pSTAT5-ir we report in this work are specifically produced by PRL, whereas other endocrine agents signalling through the Jak/STAT pathway, such as growth hormone(35), do not contribute substantially to STAT5 phosphorylation in the studied cases. This is further supported by the fact that pSTAT5 appears only in centres that show expression of PRLR in the brain of male mice (Fig 1, Fig S2), whereas the distribution of growth hormone receptor does not fit the pattern of pSTAT5-ir(35,36). In addition, the reduced pSTAT5-ir found in males not supplemented with exogenous PRL indicates very low levels of PRL signalling in basal conditions, what fits the circulating levels reported for the hormone for males(10,37). In fact, the dorsal aspect of the Arc (presumed location of the TIDA neurons as revealed by tyrosine hydroxylase immunohistochemistry, 38), displays scarce pSTAT5-ir in Male Control mice (Fig 2T).

Altogether, the male mouse brain shows little responsiveness to PRL under basal conditions, but this responsiveness would increase significantly when higher levels of the hormone reached the brain. Hence, PRL would influence the brain of males especially under some physiological conditions in which there is substantially increased hypophyseal or extra-hypophyseal PRL release, for instance during the dark period of the day (37), after mating(39) or as part of the stress response (40).

### Sexually dimorphic PRL-derived signalling in the mouse brain

In comparison to males, female mice displayed a more extensive distribution of pSTAT5-ir, mainly in the hypothalamus and basal telencephalon, but also in different nuclei of the thalamus, midbrain and brainstem. Patterns of pSTAT5-ir in males were always a completely overlapping fraction of those of female mice, with no single brain structure labelled in males but not in females. Furthermore, a number of nuclei with pSTAT5 expression in both sexes displayed significantly higher pSTAT5-ir density in females than in males (the AVPe/VMPO, Arc, DM and MePD), but none of the analysed nuclei showed higher levels of pSTAT5-ir in males (Fig 3). Altogether, our study reveals a clear, female-biased sexual dimorphism in PRL-derived signalling in the mouse brain upon exogenous PRL administration. In fact, this sexually dimorphic responsiveness to PRL would be extensive to specific functional systems of the brain, for instance to the Sociosexual Behavioural Network (SBN), the core system for the integration of social and reproductive behaviour (41). In female mice, virtually all nodes of the SBN display apparent PRL responsiveness (4), whereas males lack pSTAT5-ir or display marginal levels thereof (MePD) in all of them (the lateral septum, BST, medial amygdala or PAG, see Figs 1 and 2) except for the VMH, even after exogenous PRL administration and even if they express the PRL receptor (Fig 1, Fig S2). This suggests that, in these areas, either PRL responsiveness is regulated through different receptor activation pathways in males and females, or PRL signalling is downregulated, leading to a lower implication of PRL in the regulation of sociosexual behaviours in males. However, this does not exclude a role in some male-specific behaviours, as we will further discuss below.

### Prolactin and Testosterone: Mutual Neuroendocrine Regulation

One of the main outcomes of the present work is the identification of a putative regulatory role of testosterone in PRL signalling in the male mouse brain. Orchidectomy and consequent testosterone withdrawal led to a significant decrease in pSTAT5-ir density in the SFi, the Pa and both the central/dorsomedial and lateral VMH (Fig 4). This indicates a positive regulatory role of testosterone on PRL signalling in those brain regions. The actual (direct or indirect) mechanism of this process is not clear. Even though testosterone has documented inhibitory effects on hypophyseal PRL release in male rats(29) and mice(30), the effect of castration on pSTAT5-ir cannot be attributed to differences in circulating PRL, since intact and castrated males were supplemented with equivalent levels of exogenous PRL. In addition, this effect is not likely explained by a facilitated access of systemic PRL to the brain, as our results reveal that the positive effect of castration on pSTAT5-ir is not generalised but rather specific and restricted to discrete brain sites. Therefore, our results suggest that testosterone might upregulate PRLR expression or perhaps signalling downstream the PRLR in specific neural populations. Further research is required to elucidate the specific action of testosterone on PRL signalling in male mice.

An additional question of interest is the identity of the actual biological modulator of PRL signalling we report here. Testosterone can be locally metabolized into two different neuroactive steroids: it can be reduced to dihydrotestosterone (42), which binds to androgen receptors, or aromatised to estradiol (43) that then binds to oestrogen receptors. Our results make it very unlikely that this regulation occurs through the aromatisation of testosterone to estradiol. Although all the nuclei where PRL signalling is affected by castration express at least one form of oestrogen receptor(44), none of them displays aromatase activity in the male mouse brain(45). Conversely, all of the nuclei where PRL-derived signalling decreased with castration express androgen receptors, with the Pa and VMH showing very high levels(45). Hence, evidence suggests that the effect of testosterone on PRL signalling that we report here is likely mediated by androgen rather than oestrogen receptors.

In addition to the reported regulation of testosterone on central PRL signalling, our data, together with published evidence, suggest a reciprocal regulatory action of PRL on the hypothalamus-pituitary-gonadal (HPG) axis and ultimately on testosterone release. According to the literature, PRL is thought to upregulate the HPG axis and testosterone secretion in males, since PRL infusions stimulate the increase in serum testosterone in male rats, whereas immunological disruption of endogenous prolactin eliminates this effect (14,46). This regulatory action of PRL on the HPG axis likely occurs through the kisspeptin system, which regulates gonadotropin-releasing hormone (GnRH) expressing neurons in the brain(47) and thereby activates the HPG axis. Therefore, the expression of PRL receptors by kisspeptin neurons(48) allows for a direct role of PRL on HPG axis control and testosterone secretion. In this context, our results provide support for the hypothesis that the brain kisspeptin system is subject to PRL regulation in male mice, too, since our sample of male mice displayed substantial pSTAT5-ir in the hypothalamic nuclei containing kisspeptin-positive neurons(49), the AVPe and the Arc (Figs 2j and 2r). Nevertheless, this possibility requires experimental proof by assessing the coexpression of kisspeptin and pSTAT5 immunoreactivity in the aforementioned nuclei.

### Prolactin signalling in the male brain: functional implications

Our findings confirm that the male mouse brain is indeed responsive to PRL, implying that PRL likely exerts certain actions in the male mouse brain, provided sufficient levels of the hormone. The nature of these actions is mostly unknown, but the multiple functional studies on central PRL in female mice provide a good comparative framework to contextualize our findings.

In this respect, the most prominent (but poorly understood) function of PRL in males is the regulation of sexual behaviour (3,16). It has been shown that central action of PRL after mating suppresses sexual behaviour as part of the satiatory mechanisms operating after ejaculation (17,22). Similar inhibitory action of PRL onto sexual behaviour occurs under chronic hyperprolactinaemia(50–52). Prolactin has been proposed to do so by modulating the function of the major dopaminergic circuits in the brain, closely involved in the motor component of male sexual function (17, 53). In support of this, acute elevations of circulating PRL after copulation lead to significant decreases in dopaminergic activity in the target sites of the major dopaminergic pathways (17), including the striatum (23,54) and the MPA region (55). Several studies have shown that, indeed, PRL is capable of modulating the dopaminergic activity of these pathways at several levels (54–58). However, according to our findings, none of the nuclei originating these dopaminergic pathways (17) display PRL-derived signalling neither in female nor in male mice, even after injection of exogenous PRL (Figs 1 and 2). Furthermore, in situ hybridisation in the mouse brain does not show any expression of PRLR mRNA within these pathways, neither in their sites of origin (substantia nigra or ventral tegmental area), nor in their projection sites (caudatus putamen or nucleus accumbens) (Fig S2), although certain studies did find evidence of the receptor expression in the rat (59,60). Hence, this suggests that the proposed action of PRL over these dopaminergic systems to regulate male sexual behaviour should rather be indirect. In this vein, our study evidences a testosterone-dependent action of PRL on two nuclei of the male brain that have also been involved in male sexual behaviour regulation: the VMH and the Pa (Figs 2 and 4). The VMH is sexually dimorphic (61), known to promote the expression of copulatory behaviour in females (62) and to inhibit mounting in male rats (63), whereas the Pa contains magno- and parvocellular oxytocin (OXT) neurons involved in the peripheral and central regulation of male sexual behaviour, respectively (64–67).

The interaction between OXT and PRL at the level of the Pa can also have a role in the regulation of parental behaviours. There is solid evidence indicating that PRL (68) and OXT are fundamental in the control of maternal behaviour (69,70) and it is likely that they have an equivalent role in males. In this line, PRL facilitates pup-sensitisation in male rats (71) in a similar fashion as in females (68) and has further roles in other experimental models for paternal behaviour (72). Tachikawa and collaborators (73) showed that paternal behaviour can be induced in male mice (which are usually infanticidal) by postcopulatory cohabitation with a maternal female, a process apparently dependent on OXT release in the nucleus accumbens (Acb) (74,75). In turn, Dolen and collaborators (76) demonstrated that the OXT innervation of the Acb originates in the Pa (see also(77), which, according to our findings, is responsive to PRL in males (Fig 2M). Therefore, PRL might participate in the regulation of parental behaviour in male mice, too, by acting on Pa OXT neurons. This hypothesis requires further study.

Our results indicate a modulatory role of PRL onto the Arc and Pe nuclei of males (2R-T), both of which contain the dopaminergic neuron populations responsible for the inhibitory feedback control of hypophyseal PRL release(78): the tuberoinfundibular (TIDA) and tuberohypophyseal (THDA) neurons in the dorsomedial and rostral Arc(79), and the periventricular hypophyseal neurons (PHDA) in the Pe(80). In fact, in female rats PRL feedback signal in these dopaminergic populations is conveyed through the Jak/STAT pathway, specifically through STAT5b (81). Therefore, our results in the Arc and Pe likely reflect the (expected) occurrence of the same feedback mechanism in male mice.

Prolactin also plays an important role in the modulation of anxiety and stress. Concerning anxiety, studies in female rats indicate that PRL has anxiolytic properties when administered i.c.v. or i.v., whereas chronic i.c.v. antisense blocking of the PRLR expression results in increased anxiety(82). In contrast to females, PRL has a weak anxiolytic effect in males(82). This intersexual difference might relate to our findings of a sexually dimorphic responsiveness to PRL in the central structures directly involved in the regulation of anxiety and emotionality, the Ce and the anterodorsal BST (83). In these nuclei, males are devoid of pSTAT5-ir (Fig 1) although they show PRLR mRNA transcript (Fig 1 and fig S2), whereas females show quite abundant immunostaining, especially during pregnancy and lactation (4).

Besides being a central agent in acute stress response (11), PRL also has an important role in stress modulation. Thus, central inhibition of the expression of PRLR by i.c.v. antisense blocking results in an enhanced activation of HPA axis in male and female rats (82), demonstrating a downregulation of the stress response by PRL. The presence of pSTAT5-ir in the Pa of males (in levels equivalent to females, see Figs 2m, 2n and 3) suggests that this modulatory effect of PRL on the reactivity of the HPA axis would probably take place through PRL action on corticotropin-releasing factor (CRF) or vasopressin (AVP) cells (84). In fact, PRL is a well-documented regulator of the function of magno- and parvocellular vasopressinergic neurosecretory cells at the level of the hypothalamic Pa (85–87). Although effects of PRL on these parvocellular AVP cells has been proven to be mediated by ERK/MAPK pathway (87), our data in males and females suggest that the Jak/STAT5 signalling cascade can also be involved. This would imply a non-dimorphic regulatory role of PRL on the stress response (in addition to its dimorphic role on anxiety), which requires further research.

## Conclusions

This work characterizes the distribution of PRL responsiveness in the male mouse brain and its dependence on circulating PRL levels, sexual dimorphism and gonadal steroid regulation. Male mice display specific patterns of PRL-derived signalling only in the presence of high levels of circulating PRL. These patterns comprise mainly hypothalamic nuclei and some telencephalic sites, and are also more limited in extension and density than the equivalent patterns in female mice, evidencing a clear sexual dimorphism in favour of females. Furthermore, PRL-derived signalling in the male brain is regionally dependent on testosterone input, indicating a reciprocal regulation between both hormones. This work confirms that PRL exerts central actions in males, suggesting a possible role of the “maternal hormone” in the regulation of male sexual behaviour, PRL release feedback control or in the regulation of stress response, among others.

## ACKNOWLEDGEMENTS

Funded by the Spanish Ministry of Economy and Competitiveness-FEDER (BFU2016-77691-C2-2-P and C2-1-P) and the Generalitat Valenciana (PROMETEO/2016/076) and the *Universitat Jaume I de Castelló* (UJI-B2016-45). H.S-L has been a predoctoral fellow of the FPU (*Formación de Profesorado Universitario*) programme of the Spanish Ministry of Education, Culture and Sport.

The authors are indebted to The Allen Brain Institute for providing open access to their invaluable material and permission to use and publish its data.

## Abbreviations

3V: third ventricle
AAD: anterior amygdaloid area, dorsal part
AC/ADP: anterior commissural nucleus/anterodorsal preoptic nucleus region ac anterior commissure
Acb: nucleus accumbens
ACo: anterior cortical amygdaloid nucleus AHi amygdalohippocampal area
AHP: anterior hypothalamic area, posterior part Aq aqueduct (Sylvius)
Arc: arcuate hypothalamic nucleus AVP arginine-vasopressin
AVPe: anteroventral periventricular nucleus Bar Barrington’s nucleus
BLA: basolateral amygdaloid nucleus, anterior part BLV basolateral amygdaloid nucleus, ventral part BMP basomedial amygdaloid nucleus, posterior part
BSTIA: bed nucleus of the stria terminalis, intraamygdaloid division BSTLD bed nucleus of the stria terminalis, lateral division, dorsal part BSTLP bed nucleus of the stria terminalis, lateral division, posterior part BSTLV bed nucleus of the stria terminalis, lateral division, ventral part BSTMA bed nucleus of the stria terminalis, medial division, anterior part
BSTMV: bed nucleus of the stria terminalis, medial division, ventral part
BSTMPI: bed nucleus of the stria terminalis, medial division, posterointermediate part BSTMPL bed nucleus of the stria terminalis, medial division, posterolateral part BSTMPM bed nucleus of the stria terminalis, medial division, posteromedial part
cc: corpus callosum
Ce: central amygdala
CeC: central amygdaloid nucleus, capsular part CeL central amygdaloid nucleus, lateral division CeM central amygdaloid nucleus, medial division CPu caudate putamen (striatum)
CRF: corticotropin-releasing factor CxA cortex-amygdala transition zone D3V dorsal third ventricle
DEn: dorsal endopiriform nucleus Dk nucleus of Darkschewitsch
DLG: dorsal lateral geniculate nucleus DLPAG dorsolateral periaqueductal grey DM dorsomedial hypothalamic nucleus DMPAG dorsomedial periaqueductal grey DMTg dorsomedial tegmental area
DpG: deep grey layer of the superior colliculus DpMe deep mesencephalic nucleus
DpWh: deep white layer of the superior colliculus DR dorsal raphe nucleus
DTg: dorsal tegmental nucleus f fornix
fi: fimbria of the hippocampus
HDB: nucleus of the horizontal limb of the diagonal band I intercalated nuclei of the amygdala
ic: internal capsule
IF: interfascicular nucleus
InC: interstitial nucleus of Cajal
InG: intermediate grey layer of the superior colliculus InWh intermediate white layer of the superior colliculus
IPAC: interstitial nucleus of the posterior limb of the anterior commissure La lateral amygdaloid nucleus
LA: lateroanterior hypothalamic nucleus LC locus coeruleus
LDTg: laterodorsal tegmental nucleus LGP lateral globus pallidus
LH: lateral hypothalamic area LHb lateral habenular nucleus lo lateral olfactory tract
LOT: nucleus of the lateral olfactory tract LPAG lateral periaqueductal grey
LPBC: lateral parabrachial nucleus, central part LPBD lateral parabrachial nucleus, dorsal part LPBE lateral parabrachial nucleus, external part LPBI lateral parabrachial nucleus, internal part LPBV lateral parabrachial nucleus, ventral part LSD lateral septal nucleus, dorsal part
LSI: lateral septal nucleus, intermediate part LSV lateral septal nucleus, ventral part
LV: lateral ventricle
MCLH: magnocellular nucleus of the lateral hypothalamus mcp middle cerebellar peduncle
MCPO: magnocellular preoptic nucleus ME median eminence
MeA: medial amygdaloid nucleus, anterior part
MePD: medial amygdaloid nucleus, posterodorsal part MePV medial amygdaloid nucleus, posteroventral part MGD medial geniculate nucleus, dorsal part
MGM: medial geniculate nucleus, medial part MGP medial globus pallidus
MGV: medial geniculate nucleus, ventral part MHb medial habenular nucleus
ml: medial lemniscus
mlf: medial longitudinal fasciculus MnPO median preoptic nucleus MnR median raphe nucleus
MPA: medial preoptic area
MPB: medial parabrachial nucleus
MPBE: medial parabrachial nucleus, external part MPO medial preoptic nucleus
MS: medial septal nucleus mt mammilothalamic tract MTu medial tuberal nucleus
MZMGV: marginal zone of the medial geniculate opt optic tract
OT: oxytocin
ox: optic chiasm
Pa: paraventricular nucleus
PaLM: paraventricular nucleus, lateral magnocellular part PaMM paraventricular nucleus, medial magnocellular part PaPO paraventricular nucleus, posterior part
PaV: paraventricular nucleus, ventral part
PAG: periaqueductal greyPe periventricular hypothalamic nucleus
PeF: perifornical nucleus
PH: posterior hypothalamic area
PIL: posterior intralaminar thalamic nucleus Pir piriform cortex
PLCo: posterolateral cortical amygdaloid nucleus PnC pontine reticular nucleus, caudal part
PnO: pontine reticular nucleus, oral part PnR pontine raphe nucleus
PnV: pontine reticular nucleus, ventral part
PoT: posterior thalamic nuclear group, triangular part PP peripeduncular nucleus
PRL: prolactin
PSTh: parasubthalamic nucleus
PV: paraventricular thalamic nucleus
PVA: paraventricular thalamic nucleus, anterior part RCh retrochiasmatic area
RMg: raphe magnus nucleus
s5: sensory root of the trigeminal nerve SCh suprachiasmatic nucleus
SFi: septofimbrial nucleus SFO subfornical organ
SG: suprageniculate thalamic nucleus SI substantia innominata
SLEA: sublenticular extended amygdala sm stria medullaris of the thalamus SNC substantia nigra, pars compacta SNR substantia nigra, reticular part SO supraoptic nucleus
SPa: subparaventricular zone of the hypothalamus
st: stria terminalis
STh: subthalamic nucleus
SubCD: subcoeruleus nucleus, dorsal part SubCV subcoeruleus nucleus, ventral part SuM supramammillary nucleus
Te: terete hypothalamic nucleus Tu olfactory tubercle
VEn: ventral endopiriform nucleus VLG ventral lateral geniculate nucleus
VLGMC: ventral lateral geniculate nucleus, magnocellular part VLGPC ventral lateral geniculate nucleus, parvicellular part VLPAG ventrolateral periaqueductal grey
VLPO: ventrolateral preoptic nucleus
VMH: ventromedial hypothalamic nucleus
VMHc: ventromedial hypothalamic nucleus, central part VMHdm ventromedial hypothalamic nucleus, dorsomedial part VMHvl ventromedial hypothalamic nucleus, ventrolateral part VMPO ventromedial preoptic nucleus
VOLT: vascular organ of the lamina terminalis VP ventral pallidum
VRe: ventral reuniens thalamic nucleus vsc ventral spinocerebellar tract
VTA: ventral tegmental area ZI zona incerta

